# Histone demethylome map reveals combinatorial gene regulatory functions in embryonic stem cells

**DOI:** 10.1101/2020.08.27.269514

**Authors:** Yogesh Kumar, Pratibha Tripathi, Pushkar Dakle, Majid Mehravar, Varun K. Pandey, Michael J. Bullen, Zhongming Zhang, Dhaval Hathiwala, Marc Kerenyi, Andrew Woo, Alireza Ghamari, Alan B. Cantor, Lee H. Wong, Jonghwan Kim, Kimberly Glass, Guo-Cheng Yuan, Luca Pinello, Stuart H. Orkin, Partha Pratim Das

## Abstract

Epigenetic regulators and transcription factors establish distinct regulatory networks for gene regulation to maintain the embryonic stem cell (ESC) state. Although much has been learned regarding individual epigenetic regulators, their combinatorial functions remain elusive. Here, we report previously unknown combinatorial functions of histone demethylases (HDMs) in gene regulation of mouse ESCs. Generation of a histone demethylome (HDMome) map of 20 well-characterized HDMs based on their genome-wide binding revealed co-occupancy of HDMs in different combinations: KDM1A-KDM4B-KDM6A and JARID2-KDM2B-KDM4A-KDM4C-KDM5B largely co-occupy at enhancers and promoters, respectively. Mechanistic studies uncover that KDM1A-KDM6A combinatorially modulates P300/H3K27ac, H3K4me2 deposition and OCT4 recruitment that directs the OCT4/CORE regulatory network for target gene expression; while co-operative actions of JARID2-KDM2B-KDM4A-KDM4C-KDM5B control H2AK119ub1 and bivalent marks of polycomb-repressive complexes that facilitate the PRC regulatory network for target gene repression. Thus, combinatorial functions of HDMs differentially impact gene expression programs in mESCs.

---

The self-renewal and pluripotency are two hallmarks of embryonic stem cells (ESCs), which are established and maintained through distinct transcriptional regulatory networks ^1^. Each of these regulatory networks encompasses a set of ESC-specific transcription factors (TFs), co-factors and chromatin/epigenetic regulators to control chromatin organization and gene regulation ^2^. Comprehensive studies using the genome-wide occupancy of these factors and their links to target gene expression revealed three principal functionally distinct regulatory networks in mouse ESCs (mESCs) – CORE (active), MYC (active) and polycomb/PRC (repressive) that maintain the mESC state (i.e. self-renewal and pluripotency) ^3–6^.

Histone demethylases (HDMs) are a class of epigenetic regulators, which consists of 20 well-characterized individual members that “remove” methyl group(s) from specific lysine (K) and arginine (R) residue(s) on histone tails to modulate chromatin structure for the gene regulation ^7^. Notably, the demethylase activity of HDMs is more focused on lysine (K) than arginine (R) residues; thus, HDMs are often called lysine (K) demethylases – KDMs. HDMs are divided into two broad “classes” based on the mechanisms by which they demethylate their substrates/histone mark(s): LSD-HDMs and JMJC-HDMs ^7, 8^. LSD1/KDM1A and LSD2/KDM1B are the two members of the LSD-HDM class. The remaining HDM members belong to the JMJC-HDM class, and are further “sub-classified” based on their sequence homology, domains and substrate specificity; namely, KDM2, KDM3, KDM4, KDM5 and KDM6 ^7, 9, 10^. Several of these individual HDMs play critical roles in controlling the ESC state ^5, 11–18^. However, how HDMs function in a combinatorial manner has not been fully explored. Prior studies, including ours, have examined combinatorial functions of only selected HDMs (KDM4A, KDM4B, KDM4C) in ESCs ^5, 15^. To gain a better appreciation of the roles of HDMs, we have expanded the scope to interrogate the combinatorial functions of all 20 HDMs and how they impact gene regulatory functions in ESCs.

Here, we have constructed a histone demethylome (HDMome) map through determination of genome-wide occupancy of the HDMs in murine ESCs (mESCs). This analysis revealed that multiple HDMs share the same binding sites at enhancer and promoter regulatory regions. KDM1A-KDM4B-KDM6A largely co-occupy at enhancer regions and JARID2-KDM2B-KDM4A-KDM4C-KDM5B co-occupy at promoter regions. Comprehensive genomic analyses demonstrate that KDM1A-KDM4B-KDM6A collaborates with ESC-TFs and belongs to the CORE network (active); whereas JARID2-KDM2B-KDM4A-KDM4C-KDM5B co-operates with polycomb repressive complexes 1 and 2 (PRC1 and PRC2) and assign to the PRC network (repressive). Furthermore, KDM1A and KDM6A combinatorially modulate P300/H3K27ac, H3K4me2 deposition and OCT4 recruitment at enhancers to direct the CORE regulatory network for target gene expression. In contrast, JARID2, KDM2B, KDM4A, KDM4C and KDM5B co-operatively regulate H2AK119ub1 (of PRC1) and bivalent (H3K27me3-H3K4me3) marks (of PRC2) at promoters and enable the PRC regulatory network for target gene repression. Hence, our findings provide mechanistic insights how combinatorial actions of HDMs coordinate gene expression programs in mESCs.

## Results

### The HDMome map reveals combinatorial co-occupancy of multiple HDMs

We conducted ChIP-seq (for antibodies specific to HDM/s) and/or *in vivo* biotinylation-mediated ChIP-seq (Bio-ChIP-seq) ^5, 19^ (for Flag-Biotin (FB)-tagged HDM mESC line/s) of 20 HDMs in mESCs (Extended Data Fig. 1a and Supplementary Table 1). Of note, the majority of these HDMs are expressed in mESCs ^5^. Genome-wide occupancy data determined by ChIP-seq and/or Bio-ChIP-seq was used to construct a histone demethylome (HDMome) map (Fig. 1a,b; Extended Data Fig. 1a; Supplementary Table 1), which was utilized to assess the combinatorial co-occupancy of HDMs. Furthermore, this genome-wide binding dataset was compared with available datasets of several HDMs (KDM1A, KDM2A, KDM2B, KDM4A, KDM4C, KDM5B, KDM5C, KDM6A, KDM6B and JARID2) that demonstrated our dataset and available datasets are comparable for each of these HDMs, and they displayed significant binding enrichment (Extended Data Fig. 1a and Supplementary Table 1, 2). In addition, we generated genome-wide binding data of the remaining HDMs (KDM3A, KDM3B, KDM4B, KDM4D, KDM5A, KDM5D, PHF2, PHF8, UTY and JMJD6), which also exhibited substantial binding enrichment (Extended Data Fig. 1a and Supplementary Table 1).

**Fig. 1.**
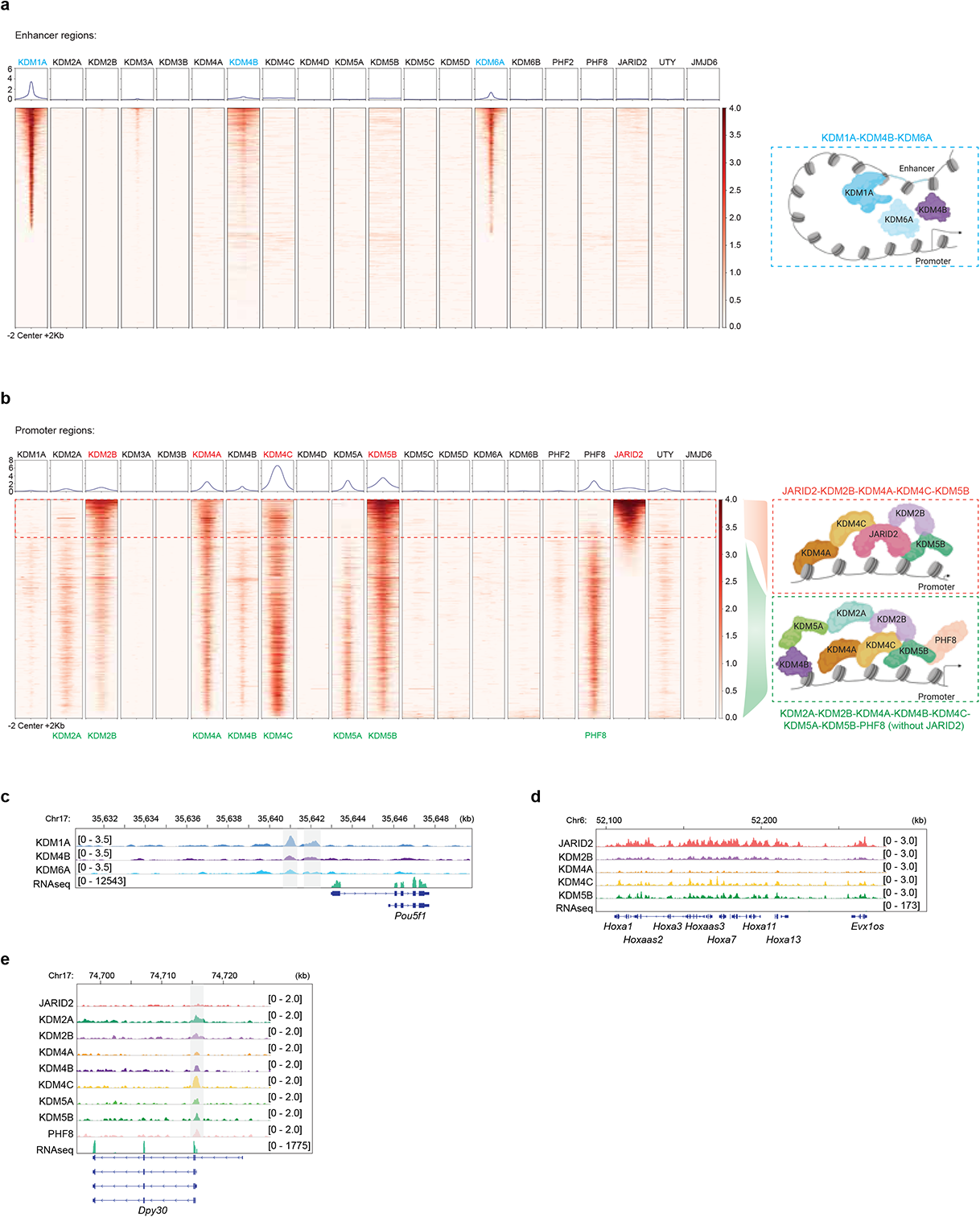
The HDMome map reveals combinatorial co-occupancy of multiple HDMs. (a) Heat map representation of co-occupancy of all HDMs at mESC-specific enhancers (8,794). Plots are centered on the region midpoint ± 2kb. Relative ChIP-seq peak intensities are indicated. The cartoon displays co-occupancy of KDM1A-KDM4B-KDM6A at enhancers. All ChIP-seq data were represented here. (b) Heat map representation of co-occupancy of all HDMs at promoters. Plots are centered on the region midpoint ± 2kb. Relative ChIP-seq peak intensities are indicated. The cartoon shows the co-occupancy of JARID2-KDM2B-KDM4A-KDM4C-KDM5B (with JARID2) and KDM2A-KDM2B-KDM4A-KDM4B-KDM4C-KDM5A-KDM5B-PHF8 (without JARID2) at promoters. All ChIP-seq data were represented here. (c,d.e) Genomic tracks illustrate ChIP-seq normalised reads for multiple HDMs (in three different combinations) at *Pou5f1* (c), *HoxA* (d) and *Dpy 30* (e) gene loci. RNA-seq tracks demonstrate expression of these genes in wild-type mESCs.

The co-occupancy (i.e. shared/ common/ overlapped binding regions) of HDMs was determined at enhancers and promoters ^20, 21^. We used 8,794 mouse ESC-specific enhancers that were identified based on co-occupancy of ESC-specific TFs (OCT4, NANOG, SOX2, KLF4, ESRRB), mediators (MED1), enhancer histone marks (H3K4me1, H3K27ac) and DNase I hypersensitivity ^21^. While ±2kb of TSS (transcription start sites) were used as promoters. Binding peaks of each HDMs (the top 5,000 peaks based on ChIP and/or Bio-ChIP-seq signal intensities) were plotted within these enhancers and promoters to assess their co-binding. We observed co-occupancy of KDM1A-KDM4B-KDM6A mainly at enhancers (Fig. 1a,c and Extended Data Fig. 1b,d); whereas JARID2-KDM2B-KDM4A-KDM4C-KDM5B and KDM2A-KDM2B-KDM4A-KDM4B-KDM4C-KDM5A-KDM5B-PHF8 (without JARID2) largely co-occupied at promoters (Fig. 1b,d,e and Extended Data Fig. 1c,e,f).

Further analyses demonstrated that KDM1A-KDM4B-KDM6A co-occupied regions mainly overlapped with active H3K27ac, H3K4me1, H3K4me2 enhancer marks, and KDM2A-KDM2B-KDM4A-KDM4B-KDM4C-KDM5A-KDM5B-PHF8 (without JARID2) co-occupied regions largely overlapped with active H3K4me3 promoter mark; both target gene sets were correlated with gene expression (Extended Data Fig. 2a,c,d and Supplementary Table 3). In contrast, JARID2-KDM2B-KDM4A-KDM4C-KDM5B co-binding regions overlapped with repressive H2AK119ub1 and bivalent (H3K27me3–H3K4me3) promoter marks, and associated genes were correlated with gene repression (Extended Data Fig. 2b,d and Supplementary Table 3). Taken together, these data suggest that HDMs act in specific combinations to achieve either gene activation or repression in mESCs.

### Members of specific HDM sub-classes act combinatorially

As KDM4 and KDM5 sub-classes represent the largest category of HDMs (KDM4 sub-class: KDM4A, 4B, 4C, 4D; KDM5 sub-class: KDM5A, 5B, 5C, 5D), we used the members of these two sub-classes to investigate their combinatorial and overlapping functions.

For KDM4 members, we first mapped unique binding sites of KDM4A, KDM4B and KDM4C, as well as their common binding sites (in different combinations) at enhancers (8,794) and promoters (±2kb of TSS) (Fig. 2a,b and Extended Data Fig. 3a,b). Next, these binding sites were correlated with several relevant active (H3K27ac, H3K4me1, H3K4me3) and repressive (H2AK119ub1, H3K27me3, H3K9me3) histone marks. KDM4D was excluded, as it does not bind significantly at enhancers or promoters (Fig. 1a,b). Enhancers encompassed mainly KDM4B unique targets, which overlapped with active H3K27ac and H3K4me1 marks and correlated with gene activation (Fig. 2a,d,g and Extended Data Fig. 3a) ^5^. On the other hand, promoter regions contained largely unique targets of KDM4A, KDM4B, KDM4C and common targets of KDM4A-4C, KDM4B-4C and KDM4A-4B-4C (Fig. 2b). KDM4A unique, KDM4C unique and KDM4A-4C common targets were co-occupied with repressive H2AK119ub1 and bivalent (H3K27me3-H3K4me3) marks, and correlated with gene repression (Fig. 2b,e,h,j and Extended Data Fig. 3b). KDM4B unique targets, as well as common targets of KDM4B-4C and KDM4A-4B-4C, largely overlapped with active H3K4me3 mark (Fig. 2b and Extended Data Fig. 3c,d). However, only KDM4B unique target genes were linked to gene expression, and target genes of other combinations did not correlate to either gene activation or repression (Fig. 2e). We failed to observe significant overlap between KDM4A, KDM4B, KDM4C targets and H3K9me3 (the substrate of KDM4 members) (Fig. 2b). These data suggest that KDM4A and KDM4C functions in gene repression and have an overlapping function, whereas KDM4B functions in gene activation. These inferences support the previous findings demonstrating the overlapping function of KDM4A and KDM4C ^15^, and combinatorial functions of KDM4B and KDM4C in mESCs ^5^.

**Fig. 2.**
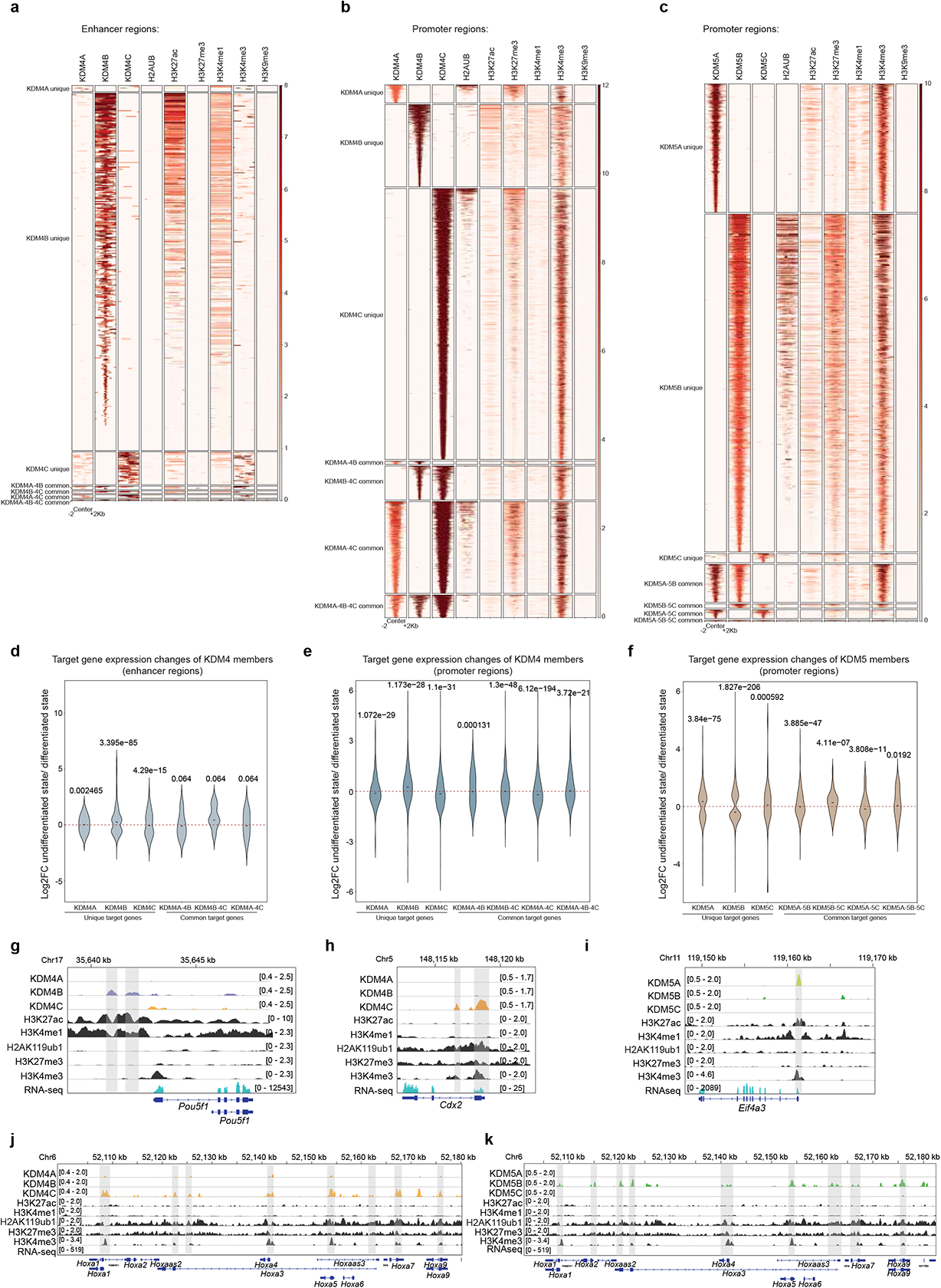
Members of specific HDM sub-classes act combinatorially. (a,b) Heat maps show binding of KDM4A, KDM4B and KDM4C along with histone marks at the ± 2kb regions around the unique and different combinations of common KDM4A, KDM4B and KDM4C binding sites within mESC-specific enhancers (a) and promoters (b). Relative ChIP-seq peak intensities are indicated. (c) A heat map displays occupancy of KDM5A, KDM5B and KDM5C and histone marks at the ± 2kb regions around the unique and different combinations of common KDM5A, KDM5B and KDM5C binding sites within promoters. Relative ChIP-seq peak intensities are indicated. (d,e) Target gene expression changes of KDM4 members (unique and different combinations of common target genes of KDM4 members) in the undifferentiated state compared to the differentiated state of mESCs at enhancers (d) and promoters (e). (f) Target gene expression changes of KDM5 members (unique and different combinations of common target genes of KDM5 members) in the undifferentiated state compared to the differentiated state of mESCs at promoters. (g,h,i,j,k) Genomic tracks represent ChIP-seq normalized reads for KDM4 and KDM5 members, as well as for histone marks at different gene loci. RNA-seq tracks demonstrate expression of these genes in wild-type mESCs.

Similar analyses were performed for KDM5 members, including KDM5A, KDM5B and KDM5C. KDM5D was excluded, as its binding was not detected at either enhancers or promoters (Fig. 1a,b). Since KDM5 members (KDM5A, KDM5B, KDM5C) binding at enhancers was infrequent (Extended Data Fig. 3e), we focused our analyses at promoters (Extended Data Fig. 3f) and found that promoters mainly included KDM5A unique, KDM5B unique and KDM5A-5B common targets (Fig. 2c). KDM5A unique targets largely overlapped with the active H3K4me3 mark and were related to gene activation (Fig. 2c,f,i). In contrast, KDM5B unique targets mainly overlapped with repressive H2AK119ub1 and bivalent (H3K27me3-H3K4me3) marks, and were connected to gene repression (Fig. 2c,f,k). Moreover, KDM5A-5B common targets predominantly co-occupied with the active H3K4me3 mark, and were linked to moderate gene expression levels (Fig. 2c,f and Extended Data Fig. 3g). Collectively, these data suggest that KDM5A and KDM5B have opposite gene regulatory functions; KDM5A associate with active gene regulatory functions, while KDM5B link to repressive gene regulatory functions.

Overall, these findings imply that KDM4 and KDM5 members execute combinatorial gene regulatory functions in mESCs.

### HDM modules co-operate with distinct ESC regulatory networks

Epigenetic regulators function with ESC-TFs to establish ESC regulatory networks for the maintenance of ESC state ^1, 2^. To reveal how HDMs collaborate with ESC-TFs and other epigenetic regulators for the gene regulation, we integrated the HDMome dataset along with binding datasets of ESC-TFs and other epigenetic regulators and mapped their binding sites at enhancers (mESC-specific 8,794 enhancers) and promoters (±2kb of TSS). The co-occupancy of multiple HDMs, ESC-TFs and other epigenetic regulators was defined as a module. Our unbiased analysis revealed three distinct modules at enhancers (named eModule – I, II, III) and promoters (named pModule – I, II, III) (Fig. 3a,b). Among these, eModule – III exhibited significant co-occupancy of KDM1A-KDM4B-KDM6A HDMs, ESC-TFs (OCT4, NANOG, SOX2), mediators (MED1,12), a cohesin complex (SMC1, NIPBL) and co-activator P300 at enhancers (Fig. 3a,c). Mediators, the cohesin complex and P300 are involved in enhancer-promoter looping at active gene loci in mESCs ^22, 23^. Conversely, pModule – I displayed significant co-occupancy of JARID2-KDM2B-KDM4A-KDM4C-KDM5B HDMs, PRC1 (RING1B, CBX7) and PRC2 (EZH2, SUZ12, JARID2) components (Fig. 3b,d). Collectively, these data hint at the possible mechanisms by which KDM1A-KDM4B-KDM6A and JARID2-KDM2B-KDM4A-KDM4C-KDM5B control gene regulation.

**Fig. 3.**
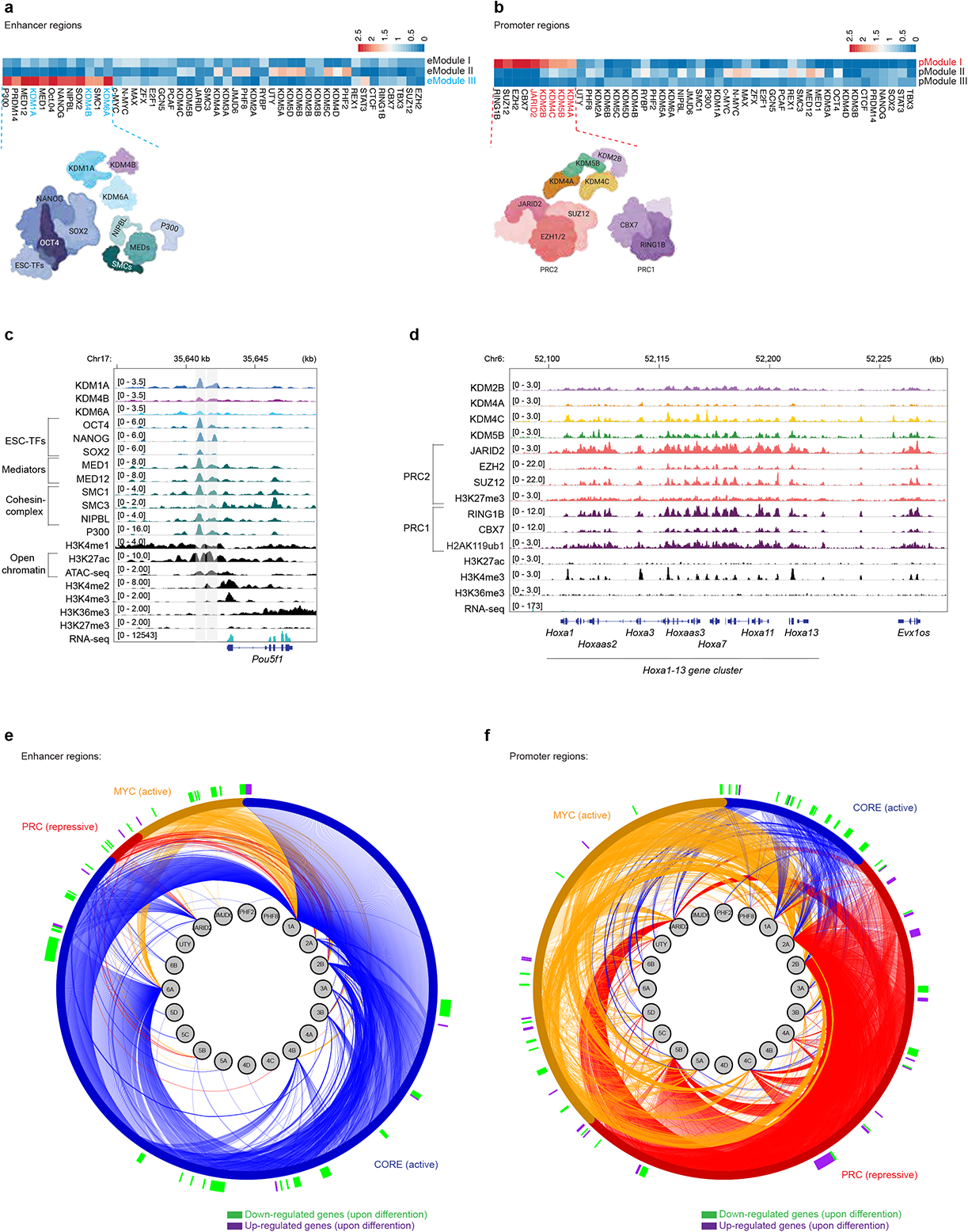
HDM modules cooperate with distinct ESC regulatory networks. (a,b) Modules are presented based on the co-occupancy of multiple HDMs, ESC-TFs and epigenetic regulators at enhancers (a) and promoters (b) in an unbiased manner. Each row represents one module; enhancer modules (eModule – I, II, III) (a), and promoter modules (pModule – I, II, III) (b) are shown. The relative binding intensities of each factors are indicated. A schematic diagram of eModule – III illustrates prominent co-occupancy of KDM1A-KDM4B-KDM6A HDMs along with ESC-TFs (OCT4, NANOG, SOX2), mediators (MED1,12), a cohesin complex (SMC1, NIPBL) and co-activator P300 at enhancers (a); whereas a schematic diagram of pModule – I illustrates significant co-occupancy of JARID2-KDM2B-KDM4A-KDM4C-KDM5B HDMs, PRC1 (RING1B, CBX7) and PRC2 (EZH2, SUZ12, JARID2) components at promoters (b). (c) The genome browser displays significant co-occupancy of KDM1A, KDM4B, KDM6A, ESC-TFs (OCT4, NANOG, SOX2), mediators (MED1, 12), the cohesin complex (SMC1/3, NIPBL), P300 and enhancer histone marks (H3K4me1, me2 and H3K27ac) at the *Pou5f1/Oct4* gene locus. ATAC-seq track is shown; both ATAC-seq and H3K27ac tracks represent open chromatin regions. The RNA-seq track exhibits expression of the *Pou5f1/Oct4* in wild-type mESCs. Highlighted regions depicted as enhancers. (d) The genome browser presents significant co-occupancy of JARID2, KDM2B, KDM4A, KDM4C, KDM5B, PRC1 (CBX7, RING1B) and PRC2 (EZH2, SUZ12, JARID2) components, as well as H2AK119ub1 (of PRC1) and H3K27me3–H3K4me3 bivalent (of PRC2) histone marks at the *HoxA* gene cluster. The RNA-seq track illustrate repression of *HoxA* cluster genes in wild-type mESCs. (e-f) HDMome–ESC regulatory networks at enhancers (e) and promoters (f). The inner ring contains all 20 HDMs; while the outer ring represents the CORE (active), MYC (active) and PRC (repressive) networks. The lines extending from these HDMs to the networks represent the binding sites of HDMs to the networks. Ticks along the exterior of the outer ring shows differentially expressed HDM bound network genes in the differentiated state compared to undifferentiated state of mESCs. Magenta tick marks indicate increased expression of HDM bound network genes upon differentiation, while green tick marks indicate decreased expression of HDM bound network genes upon differentiation.

Furthermore, we examined whether HDMs interact with existing ESC regulatory networks ^5, 6, 24^. Upon integration of HDMome, ESC regulatory networks and gene expression (undifferentiated vs. differentiated mESCs) datasets we observed that KDM1A, KDM4B and KDM6A were primarily associated with the CORE (active) network at enhancers (Fig. 3e). The majority of the KDM1A, KDM4B, KDM6A bound CORE network genes were highly expressed in the undifferentiated state and down-regulated upon mESCs differentiation (Fig. 3e). In contrast, JARID2, KDM2B, KDM4A, KDM4C and KDM5B were predominantly connected to the PRC (repressive) network at promoters (Fig. 3f). The PRC target genes bound by these HDMs were mostly repressed in the undifferentiated state and up-regulated upon mESCs differentiation (Fig. 3f). Additionally, we observed connections between individual HDMs and ESC regulatory networks (Fig. 3e,f), which indicate that individual HDMs may also act through distinct gene regulatory network(s).

These data collectively suggest that a specific combination of HDMs function with ESC-TFs and other epigenetic regulators to constitute a distinct module, which co-operates with distinct ESC gene regulatory network(s) either for gene expression or repression.

### KDM1A and KDM6A combinatorially modulate P300/H3K27ac, H3K4me2 deposition and OCT4 recruitment at enhancers

Although KDM1A-KDM4B-KDM6A co-occupy at enhancers, the co-occupancy is most striking between KDM1A and KDM6A (Fig. 1a and Extended Data Fig. 1d). In addition, compared to KDM4B targets, the KDM1A and KDM6A targets largely overlap with the CORE network (Fig. 3e). Based on these criteria, we focused analysis on KDM1A and KDM6A.

We created individual *knockout (KO)* and *double knockout (DKO)* mESC lines of *Kdm1a* and *Kdm6a* (Extended Data Fig. 4b,c,d; Supplementary Table 4). ChIP-seq data of relevant histone marks related to KDM1A and KDM6A were generated from wild-type and *KO* lines to capture changes in histone marks at “KDM1A-KDM6A co-binding sites” within enhancers (Fig. 4b and Extended Data Fig. 4h). High-confidence KDM1A-KDM6A co-binding sites were assigned within all mESC-specific enhancers (8,794) and “active” enhancers (4,346) (Fig. 4a and Extended Data Fig. 4e,f,g; Supplementary Table 5). Active enhancers comprised of H3K27ac and H3K4me1 marks but were devoid of H3K4me3 and ± 2kb of TSS as promoter regions (i.e. without ± 2kb TSS, NO H3K4me3, +H3K27ac, +H3K4me1) among all enhancers (Extended Data Fig. 4f; Supplementary Table 5). Hence, these active enhancers represent a subset of all enhancers. We obtained a total of 3,147 and 747 high-confidence KDM1A-KDM6A co-occupied regions within all (8,794) and active (4,346) enhancers, respectively (Fig. 4a and Extended Data Fig. 4g; Supplementary Table 5).

**Fig. 4.**
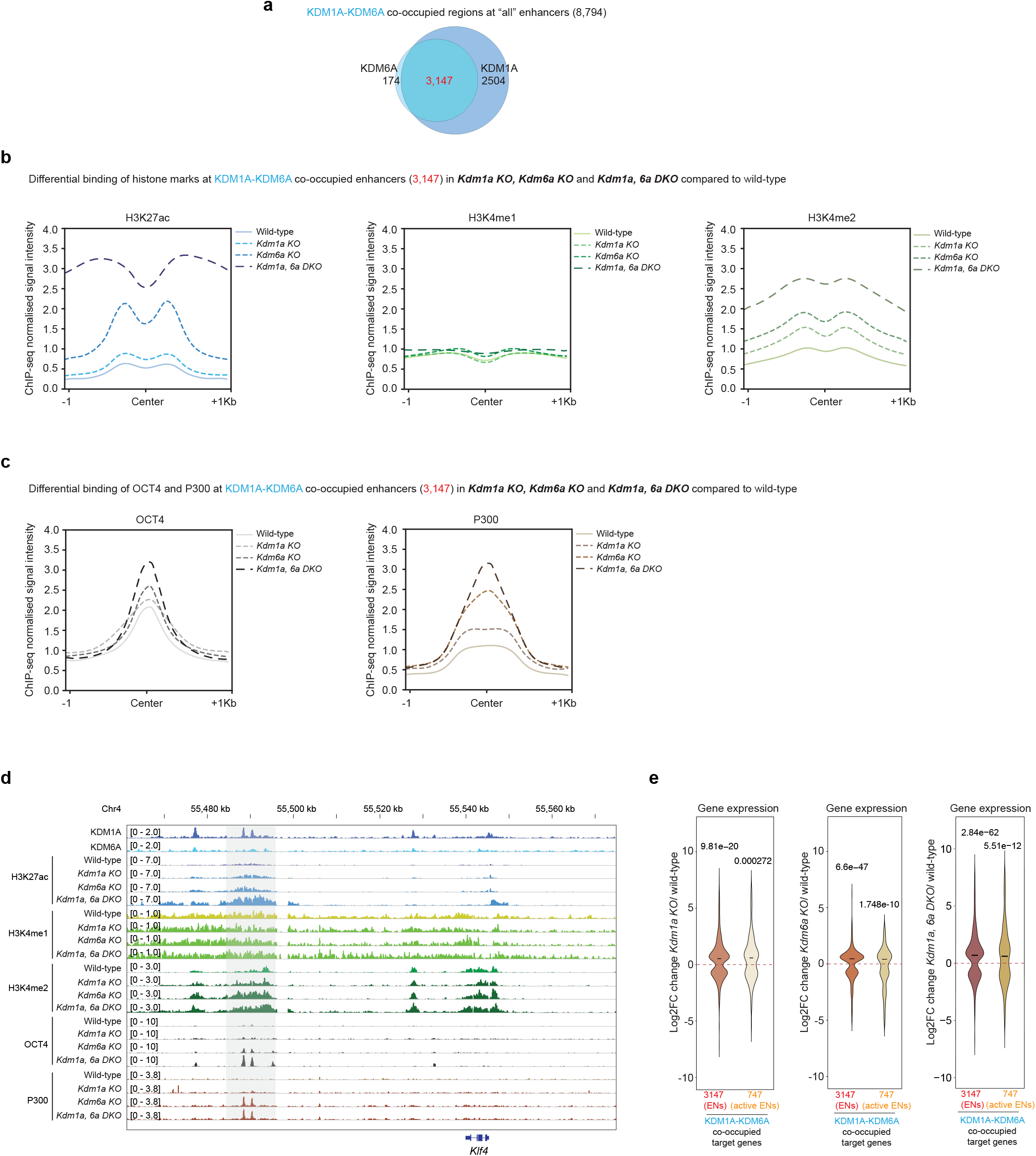
KDM1A and KDM6A combinatorially modulate P300/H3K27ac, H3K4me2 deposition and OCT4 recruitment at enhancers. (a) Venn diagram represents high-confidence KDM1A-KDM6A co-occupied sites (3,147) at all mESC-specific enhancers (ENs) (8,794). KDM1A and KDM1A-FB overlapping sites (5,651); KDM6A and KDM6A-FB overlapping sites (3,321) at all enhancers (8,794) were used to determine the KDM1A-KDM6A co-occupied sites (see Extended Data Fig. 4e). (b,c) Profile plots exhibit differential binding of H3K7ac, H3K4me1, H3K4me2 (b), and OCT4, P300 (c) at KDM1A-KDM6A co-occupied enhancers (3,147) in the *Kdm1a KO, Kdm6a KO* and *Kdm1a, 6a DKO* compared to wild-type. (d) Genomic tracks demonstrate the co-occupancy of KDM1A and KDM6A. Additionally, differential binding of histone marks (H3K7ac, H3K4me1, H3K4me2), P300 and OCT4 at KDM1A-KDM6A co-occupied enhancers from the *Kdm1a KO, Kdm6a KO* and *Kdm1a, 6a DKO* and wild-type is shown at the *Klf4* gene locus. The highlighted region displays KDM1A-KDM6A co-occupied enhancers. (e) Gene expression changes of KDM1A-KDM6A co-occupied target genes (3,147 and 747) in the *Kdm1a KO, Kdm6a KO* and *Kdm1a, 6a DKO* compared to wild-type.

Further analyses demonstrated marked increases of H3K27ac and H3K4me2 at KDM1A-KDM6A co-occupied enhancers (3,147 and 747) in *Kdm1a KO, Kdm6a KO* and *Kdm1a, 6a DKO* compared to wild-type mESCs (Fig. 4b,d and Extended Data Fig. 4h). However, the H3K4me1 mark remained unchanged at KDM1A-KDM6A co-occupied enhancers in all *KOs* (Fig. 4b,d and Extended Data Fig. 4h). The occupancy of P300 (that deposits H3K27ac) and OCT4 (a key ESC-TF, and a central player in the CORE network) was significantly increased at KDM1A-KDM6A co-occupied enhancers in *Kdm1a KO, Kdm6a KO* and *Kdm1a, 6a DKO* as well (Fig. 4c,d and Extended Data Fig. 4i). In particular, we observed increased binding or occupancy of H3K27ac, H3K4me2, P300 and OCT4 at KDM1A-KDM6A co-occupied enhancers in *Kdm1a, 6a DKO* compared to individual *Kdm1a* and *Kdm6a KOs* (Fig. 4b,c,d and Extended Data Fig. 4h,i). These comprehensive analyses indicate that KDM1A/KDM6A modulates H3K27ac through P300, while only KDM1A modulates H3K4me2 (as KDM1A is a demethylase for H3K4me1, me2) at targeted enhancers. Of note, we did not observe significant changes in H3K27me3 (a known substrate of KDM6A demethylase) at all mESC-specific enhancers (8,794), as well as at KDM1A-KDM6A co-occupied enhancers (3,147 and 747) in *Kdm6a KO* (Extended Data Fig. 4j,k), confirming the potential non-catalytic activity of KDM6A for H3K27me3 in mESCs.

Gene expression analysis demonstrated upregulation of KDM1A-KDM6A co-occupied target genes in *Kdm1a KO, Kdm6a KO* and *Kdm1a, 6a DKO* (Fig. 4e). The upregulation of KDM1A-KDM6A co-occupied target genes was positively correlated with increased binding of P300/H3K27ac and H3K4me2 at co-occupied sites in the absence of *Kdm1a* or *Kdm6a* and both (Fig. 4b,c,d,e and Extended Data Fig. 4h,i), suggesting that KDM1A and/or KDM6A represses enhancer functions. This conclusion is compatible with a report demonstrating that KDM1A/LSD1 is involved in the repression of active genes through enhancer-binding in mESCs ^25^.

Taken together, these data suggest that KDM1A and KDM6A combinatorially modulate P300/H3K27ac, H3K4me2 deposition and OCT4 recruitment at enhancers for target gene expression.

### JARID2, KDM2B, KDM4A, KDM4C and KDM5B co-operatively control H2AK119ub1 (of PRC1) and bivalent marks (of PRC2) at promoters for target gene repression

We observed a strong correlation between JARID2-KDM2B-KDM4A-KDM4C-KDM5B HDMs and the PRC network (Fig. 3b,f). As JARID2-KDM2B-KDM4A-KDM4C-KDM5B HDMs co-occupy with PRC1/H2AK119ub1 and PRC2/bivalent marks (Fig. 3b,d,f), these HDMs co-occupied targets serve as polycomb targets. To interrogate potential combinatorial mechanisms of this set of HDMs in PRC-mediated gene repression, we generated individual and combined *knockout (KO)* or knockdown (KD) mESC lines of these HDMs (Extended Data Fig. 5a,b,c) ^5, 26^.

Individual *Jarid2 KO*, Kdm2b KD, Kdm4a KD, Kdm4c KD and Kdm5b KD mESCs demonstrated significant reduction of H2AK119ub1 of PRC1, and alteration of bivalent marks (i.e. reduction of H3K27me3 and elevation of H3K4me3) of PRC2 at JARID2-KDM2B-KDM4A-KDM4C-KDM5B co-occupied promoters (Fig. 5a,b,c,d,e,f and Extended Data Fig. 5d,e,f; Supplementary Table 3). In addition, combined *Jarid2 KO*-Kdm4a KD-Kdm5b KD triple *KO*-KD (*J2*-4a-5b T*KO*-KD) mESCs also exhibited a reduction of H2AK119ub1 (of PRC1) and alteration of bivalent marks (of PRC2) at JARID2-KDM2B-KDM4A-KDM4C-KDM5B co-occupied promoters, but to a greater extent compared to individual *KO* or KD mESCs (Fig. 5a,b,c,d,e,f and Extended Data Fig. 5d,e,f). However, no significant changes of H3K9me3 and H3K36me3 were observed at JARID2-KDM2B-KDM4A-KDM4C-KDM5B co-occupied promoters in Kdm4a KD and Kdm4c KD mESCs (Extended Data Fig. 5g,h), even though both KDM4A and KDM4C exhibit H3K9me2/me3 and H3K36me2/me3 demethylase activity ^27^. Additionally, we did not detect changes in H3K36me2 at JARID2-KDM2B-KDM4A-KDM4C-KDM5B co-occupied promoters in the absence of Kdm2b (Extended Data Fig. 5i), although KDM2B is an H3K36me2, me1 demethylase ^27^.

**Fig. 5.**
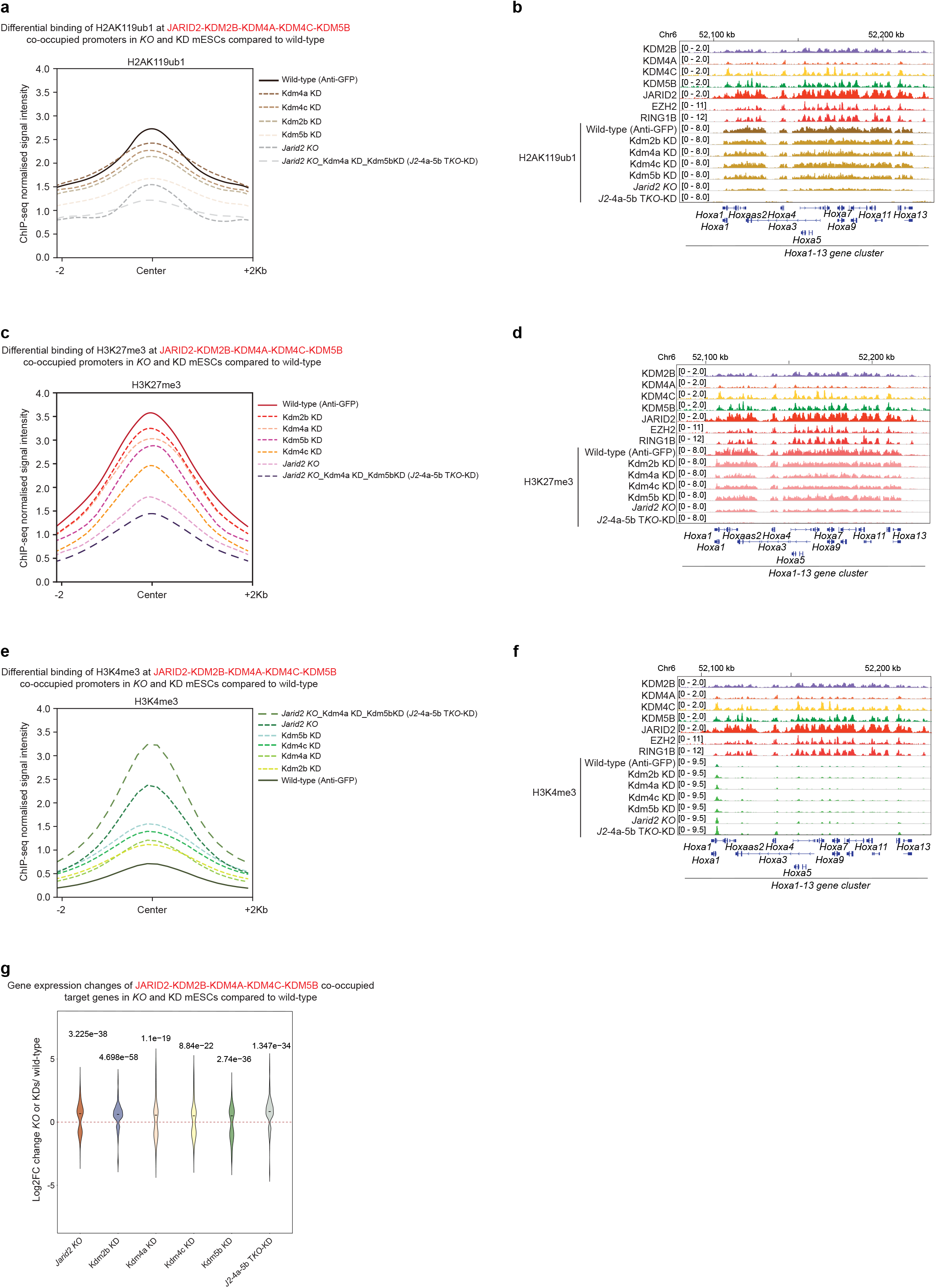
JARID2, KDM2B, KDM4A, KDM4C and KDM5B co-operatively control H2AK119ub1 (of PRC1) and bivalent marks (of PRC2) at promoters for target gene repression. (a, c, e) Profile plots represent differential binding of H2AK119ub1 (a), H3K27me3 (c) and H3K4me3 (e) at JARID2-KDM2B-KDM4A-KDM4C-KDM5B co-occupied promoters (417) in *KO* and KD mESCs compared to wild-type. (b, d, f) Genomic tracks demonstrate differential binding of H2AK119ub1 (b), H3K27me3 (d) and H3K4me3 (f) at JARID2-KDM2B-KDM4A-KDM4C-KDM5B co-occupied promoters of the *Hoxa1-13 gene cluster* in *KO* and KD mESCs compared to wild-type. (g) Gene expression changes of JARID2-KDM2B-KDM4A-KDM4C-KDM5B co-occupied target genes in *Jarid2 KO*, Kdm2b KD, Kdm4a KD, Kdm4c KD, Kdm5b KD and *Jarid2 KO*-Kdm4a KD-Kdm5b KD triple *KO*-KD (*J2*-4a-5b T*KO*-KD) mESC lines compared to wild-type.

Furthermore, gene expression analysis revealed up-regulation/de-repression of JARID2-KDM2B-KDM4A-KDM4C-KDM5B co-occupied target genes in the absence of Jarid2, Kdm2b, Kdm4a, Kdm4c and Kdm5b – individually and in combination (Fig. 5g). Taken together, these data suggest that JARID2, KDM2B, KDM4A, KDM4C and KDM5B co-operatively regulate H2AK119ub1 (of PRC1) and bivalent marks (of PRC2) at promoters for repression of their target genes.

## Discussion

Here we constructed the first comprehensive HDMome map of all 20 HDMs based on their genome-wide binding in mESCs (Fig. 1) in an effort to elucidate combinational actions of HDMs in gene regulation. The HDMome map is highly relevant in defining the targets of individual HDMs and shared targets of multiple HDMs, along with ESC-TFs, other epigenetic regulators, and histone marks (Fig. 1, 3). These analyses provide a comprehensive view of potential combinatorial actions of multiple HDMs (beyond their sub-class criteria) in controlling the turnover of histone marks *in situ* for fine-tuning gene expression programs. Following correlative genomics, we assessed how specific sets of HDMs, such as, KDM1A-KDM6A and JARID2-KDM2B-KDM4A-KDM4C-KDM5B combinatorially regulate gene expressions (Fig. 4, 5).

KDM1A/LSD1 is an H3K4me1, me2– demethylase ^8^, and co-occupies with the repressive NuRD (HDAC1/2, MBD3, Mi-2ß) complex, as well as with active ESC-TFs (OCT4, NANOG, SOX2), co-activators (mediators) and P300 (HAT) at enhancers of active ESC genes. However, H3K4me1, me2– demethylase activity of KDM1A is suppressed due to the presence of acetylated histones (H3K27ac)/HAT P300 that ultimately drives active ESC enhancer functions ^12, 28^. This proposed model has not been fully validated. Our findings are consistent with suppression of H3K4me1, me2– demethylase activity (mostly H3K4me2) of KDM1A/LSD1 at active enhancers (Fig. 4). Additionally, we inferred that KDM1A/LSD1 also regulate H3K27ac through P300 at these enhancers. Likewise, KDM6A/UTX regulate H3K27ac (via P300) and H3K4me2 (through KDM1A) at enhancers (Fig. 4). These activities of KDM6A are uncommon (in mESCs) compared to its usual activities; typically, KDM6A demethylates H3K27me2, me3 repressive marks at promoters ^7, 29^. We did not observe co-occupancy between KDM6A and H3K27me3, nor the catalytic activity of KDM6A for H3K27me3 in mESCs (Extended Data Fig. 4), as reported previously ^30^. The same study also described reciprocal regulation between KDM6A-P300 mediated H3K27ac and KDM6A-MLL4 mediated H3K4me1 at enhancers, which established a KDM6A-P300-MLL4 network at enhancers ^30^. Nonetheless, our data reveal that KDM1A/KDM6A-P300 mediate H3K27ac and KDM6A-KDM1A facilitate H3K4me2, which mutually recruits OCT4 at KDM1A-KDM6A co-occupied enhancers for gene expression (Fig. 4 and Extended Data Fig. 4). Hence, we propose a “KDM1A/KDM6A-P300-OCT4 network” at enhancers for KDM1A/KDM6A target gene expression, where KDM1A and KDM6A participate combinatorially (Fig. 6). This network may be critical for maintaining a fine-balance between acetylated (H3K27ac) and methylated (H3K4me1, me2) histones, which preserves active enhancer functions in undifferentiated mESCs.

**Fig. 6.**
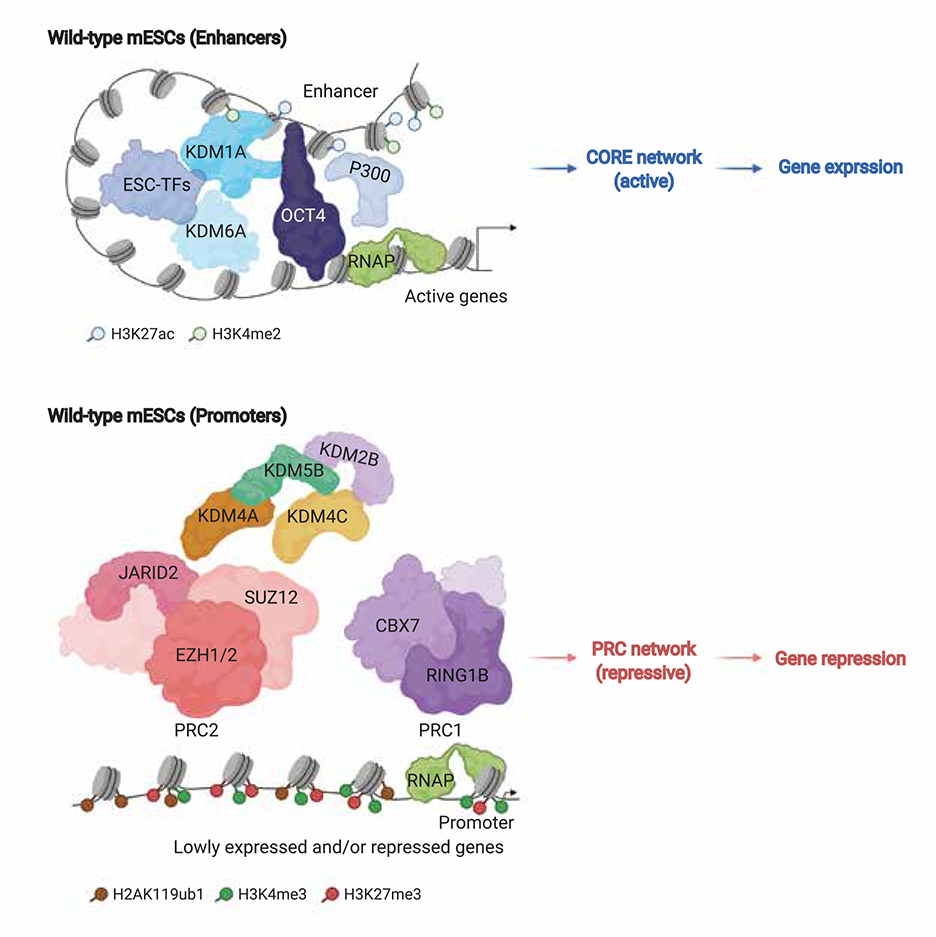
The combinatorial gene regulatory functions of HDMs in mESCs. The proposed model demonstrates that multiple HDMs function together in different combinations either at enhancers or promoters. KDM1A and KDM6A combinatorially modulate mainly P300/H3K27ac, H3K4me2 deposition and recruitment of OCT4 at enhancers to activate the CORE regulatory network for target gene expression. Conversely, JARID2, KDM2B, KDM4A, KDM4C and KDM5B co-operatively regulate H2AK119ub1 (of PRC1) and bivalent (H3K27me3-H3K4me3) marks (of PRC2) at promoters that facilitates the PRC regulatory network for target gene repression. Hence, different combinations of HDMs regulate distinct ESC gene regulatory networks in mESCs.

JARID2-KDM2B-KDM4A-KDM4C-KDM5B co-occupy with components of PRC1/H2AK119ub1 and PRC2/bivalent marks, thereby establishing a connection with the PRC gene regulatory network (Fig. 3 and Extended Data Fig. 2). Our data provided evidence that JARID2 is a critical factor (an HDM but lacks demethylase activity ^26, 31, 32^ and a component of PRC2.2 complex ^33, 34^) for the modulation of H2AK119ub1 of PRC1 and bivalent marks (H3K27me3-H3K4me3) of PRC2 at JARID2-KDM2B-KDM4A-KDM4C-KDM5B co-occupied promoters (Fig. 5). Recent studies support these conclusions and the importance of JARID2 in the PRC1-H2AK119ub1-JARID2-PRC2.2 pathway, where JARID2 acts a missing link between the PRC1 and PRC2 complexes ^35, 36^. Moreover, our work revealed that KDM2B, KDM4A, KDM4C and KDM5B HDMs principally regulate H2AK119ub1 (of PRC1) and bivalent marks (of PRC2) at JARID2-KDM2B-KDM4A-KDM4C-KDM5B co-occupied promoters as well (Fig. 5), suggesting a functional link between KDM2B, KDM4A, KDM4C, KDM5B HDMs and both PRC1 and 2 complexes. This observation is compatible with prior data, as – i) KDM2B recruits PRC1 at unmethylated CpG islands for deposition of H2AK119ub1 and gene repression in mESCs ^37^; ii) KDM4C correlates with PRC2 and gene repression in mESCs ^5^; iii) KDM5B exhibits regulation of H3K4me2, me3 at the bivalent promoters in mESCs ^38^. Moreover, the combined effect of JARID2, KDM4A and KDM5B was observed on H2AK119ub1 and bivalent marks at promoters (Fig. 5). Our data establish a critical link between JARID2, KDM2B, KDM4A, KDM4C, KDM5B HDMs (in a combinatorial fashion) and PRC1/H2AK119ub1-PRC2/(H3K27me3-H3K4me3) pathway for gene repression (Fig. 5, 6).

The HDMome atlas we have generated reveals unique combinatorial functions of HDMs that drive and orchestrate gene expression programs in mESCs.

## Methods

### Mouse embryonic stem cells (mESCs)

Mouse ESCs (mESCs) were cultured in mouse ESC media that contains DMEM (Dulbecco’s modified Eagle’s medium) (Thermo Fisher Scientific) supplemented with 15% fetal calf serum (FCS) (Thermo Fisher Scientific), 0.1mM b-mercaptoethanol (Sigma-Aldrich), 2mM L-glutamine (Thermo Fisher Scientific), 0.1mM nonessential amino acid (Thermo Fisher Scientific), 1% of nucleoside mix (Merck Millipore), 1000 U/ml recombinant leukemia inhibitory factor (LIF/ESGRO) (Merck Millipore), and 50U/ml Penicillin/Streptomycin (Thermo Fisher Scientific). mESCs were cultured at 37^°^C, 5% CO2.

### Human embryonic kidney cells (HEK293T cells)

HEK293T cells were cultured with DMEM supplemented with 10% fetal bovine serum (FBS) (Thermo Fisher Scientific) and 2% penicillin-streptomycin (Thermo Fisher Scientific). These cells were cultured at 37^°^C, 5% CO2. HEK293T cells were used only for lentiviral production.

### Cell lines

J1 mESCs (wild-type), *Kdm1a KO, Kdm6a KO* and *Kdm1a, 6a DKO* mESCs.

*Jarid2 KO* mESCs ^26, 39^.

Kdm2b KD, Kdm4a KD, Kdm5b KD mESCs; Kdm4c KD cells ^5^.

BirA, JARID2-FB, KDM1A-FB, and KDM6A-FB mESC lines.

HEK293T cells.

### Antibodies

Anti-KDM1A: Abcam ab17721 (ChIP); Santa Cruz Biotechnology sc-271720 (WB)

Anti-KDM2A: Novus Biologicals NB100-74602

Anti-KDM2B: Merck #17-10264

Anti-KDM3A: Bethyl laboratories A301-538A

Anti-KDM3B: Merck #07-1535

Anti-KDM4A: Cell Signaling #3393

Anti-KDM4B: Bethyl laboratories A301-478A

Anti-KDM4C: Abcam ab85454 (WB); Novus Biologicals NBP1-49600 (WB)

Anti-KDM4D: Santa Cruz Biotechnology sc-393750

Anti-KDM5A: Abcam ab70892

Anti-KDM5B: Bethyl laboratories A301-813A

Anti-KDM5C: Bethyl laboratories A301-034A

Anti-KDM5D: Merck #ABE203

Anti-KDM6A: Bethyl laboratories A302-374A

Anti-UTY: Abcam ab91236

Anti-KDM6B: Abcam ab95113

Anti-PHF2: Cell Signaling #3497

Anti-PHF8: Bethyl laboratories A301-772A

Anti-JMJD6: Santa Cruz Biotechnology sc-28348

Anti-JARID2: Cell Signaling #13594; Novus Biologicals NB100-2214 – (ChIP, WB)

Anti-EZH2: Cell Signaling #5246

Anti-SUZ12: Active motif # 39357; Abcam ab12073

Anti-RING1B: Bethyl laboratories A302-869A

Anti-KLF4: Abcam ab106629

Anti-KLF5: Abcam ab137676

Anti-SOX2: Abcam ab59776

Anti-OCT4: Santa Cruz Biotechnology sc-5279 (ChIP); Abcam ab19857 (WB)

Anti-P300: Abcam ab54984 (ChIP); Thermo scientific #33-7600 (ChIP)

Anti-H3K4me1: Abcam ab8895

Anti-H3K4me2: Abcam ab32356

Anti-H3K4me3: Merck #07-473

Anti-H3K9me3: Abcam ab8898

Anti-H3K27me3: Abcam ab6002

Anti-H3K27ac: Abcam ab4729

Anti-H3K36me3: Abcam ab9050

Anti-H2AK119ub1: Cell Signaling #8240

### Commercial assays

RNeasy Mini Kit (74106, Qiagen)

Ribosomal RNA depletion kit (rRNA Depletion Kit, E6310L, NEB)

NEBNext Ultra Directional RNA Library Prep Kit (E7420L, NEB)

NEBNext ChIP-seq Library Prep Master Mix Set for Illumina (E6240L, NEB)

NEBNext Ultra DNA Library Prep Kit for Illumina (E7370L, NEB)

Dynabeads MyOne Streptavidin T1 beads (65601, Thermo Fisher Scientific)

### ChIP-seq

Chromatin Immunoprecipitation (ChIP) was performed as described previously ^5, 19^. For bioChIP reactions, streptavidin beads (Dynabeads MyOne Streptavidin T1-Thermo Fisher Scientific) were used for the precipitation of chromatin, and 2% SDS was applied for the first washing step. All other steps were the same as conventional ChIP protocol. BirA expressing J1 mESCs were used as a control.

Conventional ChIP reactions were performed as described previously ^5, 39^. Input genomic DNA was used for the reference sample. Briefly, cells were trypsinized from 15cm dishes, washed twice with 1XPBS, and cross-linked with 37% formaldehyde solution (Calbiochem) to a final concentration of 1% for 8 min at room temperature with gentle shaking. The reaction was quenched by adding 2.5M glycine to a final concentration of 0.125M. Cells were washed twice with 1XPBS, and the cell pellet was resuspended in SDS-ChIP buffer (20 mM Tris-HCl pH 8.0, 150mM NaCl, 2mM EDTA, 0.1% SDS, 1% Triton X-100 and protease inhibitor), and chromatin was sonicated to around 200-500 bp. Sonicated chromatin was incubated with 5∼10µg of antibody overnight at 4°C. After overnight incubation, protein A/G Dynabeads magnetic beads (Thermo Fisher Scientific), were added to the ChIP reactions and incubated for 2-3 hours at 4°C to immunoprecipitate chromatin. Subsequently, beads were washed twice with 1 ml of low salt wash buffer (50mM HEPES pH 7.5, 150mM NaCl, 1mM EDTA, 1% Triton X-100, 0.1% sodium deoxycholate), once with 1ml of high salt wash buffer (50mM HEPES pH 7.5, 500mM NaCl, 1mM EDTA, 1% Triton X-100, 0.1% sodium deoxycholate), once with 1ml of LiCl wash buffer (10mM Tris-HCl pH 8.0, 1mM EDTA, 0.5% sodium deoxycholate, 0.5% NP-40, 250 mM LiCl), and twice with 1ml of TE buffer (10mM Tris-HCl pH 8.0, 1mM EDTA, pH 8.0). The chromatin was eluted and reverse-crosslinked simultaneously in 300µl of SDS elution buffer (1% SDS, 10mM EDTA, 50mM Tris-HCl, pH 8.0) at 65°C overnight. The next day, an equal volume of TE was added (300µl). ChIP DNA was treated with 1µl of RNaseA (10mg/ml) for 1hr, and with 3µl of proteinase K (20mg/ml) for 3hrs at 37°C, and purified using phenol-chloroform extraction, followed by QIAquick PCR purification spin columns (Qiagen). Finally, ChIP-DNA was eluted from the column with 40µl of water. For several factors, we used multiple ChIPs. In the end, all eluted ChIP-DNA samples were pooled and precipitated to enrich the ChIP-DNA material to make the libraries for high-throughput sequencing. Input ChIP samples were reserved before adding the antibodies, and these input samples were processed from reverse cross-link step until the end, same as other ChIP samples.

2-10 ng of purified ChIP DNA was used to prepare sequencing libraries, using NEBNext ChIP-seq Library Prep Master Mix Set for Illumina (NEB) and NEBNext Ultra DNA Library Prep Kit for Illumina (NEB) according to the manufacturer’s instructions. All libraries were checked through a Bio-analyser for quality control purpose. ChIP-sequencing (50bp SE reads or 150bp PE reads) was performed using Hiseq-2500 or Hiseq-4000 (Illumina).

### Generation of Flag-Biotin (FB) tagged mESC lines

Open reading frames (ORFs) of HDM genes of interest were synthesized (IDT), and cloned into BamHI-digested pEF1a-Flagbio(FB)-puromycin vector, using Gibson Assembly Master Mix (NEB). Positive FB constructs were analysed by Sanger sequencing. 10µg of each of the HDM-FB construct was electroporated into 5×10^6^ J1 wild-type mESCs, which constitutively express BirA ligase (neomycin). The electroporated cells were plated on a 15cm dish, with mESC media. After ∼24hrs, media was replaced with fresh mESC media containing 1μg/ml of puromycin (Sigma-Aldrich) and 1μg/ml of neomycin (Sigma-Aldrich), and cells were selected for 4-5days. Individual mESC colonies were picked, expanded, and tested by western blot using either streptavidin-HRP antibody (GE healthcare) (dilution 1:2000 in 5% BSA) or specific antibodies against HDM to detect the HDM-FB lines.

### Co-Immunoprecipitation

For each immunoprecipitation (IP), HDM-FB mESCs were harvested from 15 cm dish and washed twice with ice-cold PBS. The cell pellet was allowed to swell in twice the volume of Hypotonic solution (10mM HEPES-pH 7.3, 1.5mM MgCl2, 10mM KCl, 1mM DTT, 1mM PMSF and protease inhibitors), and passed through 26^1/2^-gauge needle 5 times, followed by quick centrifuge at 14000 rpm for 20-30 sec. The cloudy supernatant cytoplasmic fraction was removed, and the cell pellet was resuspended in the same volume of High salt buffer (20mM HEPES-pH 7.3, 1.5mM MgCl2, 420mM KCl, 0.2mM EDTA, 30% Glycerol, 1mM DTT, 1mM PMSF and protease inhibitors) and rotated for 1-2 hour at cold room. Then, Neutralizing buffer “without salt” (20mM HEPES-pH 7.3, 0.2mM EDTA, 20% Glycerol, 1mM DTT, 1mM PMSF and protease inhibitors) was added to the nuclear extract (NE) to bring the salt concentration to around 150mM. NE was centrifuged at 14000 rpm for 20 min at 4°C, and the supernatant was collected. The volume of supernatant was increased up to 1ml using IP buffer (combining High salt buffer and Neutralizing buffer to make final conc. of 150mM, with 1mM DTT, 1mM PMSF and protease inhibitors). Percentage of supernatant was collected as an “Input”, and rest of the supernatant was incubated with streptavidin Dynabeads (Thermo Fisher Scientific) for 2hrs at 4°C. Subsequently, immunoprecipitated protein-beads were washed 3 times with IP buffer, each for 5 minutes at 4°C. IP-ed protein and its interacting partners were eluted from the beads in 2X XT buffer (Bio-Rad) by boiling for 10 minutes; resolved on a 4-12% gradient Bis-Tris gel (Bio-Rad) and analyzed by western blot using specific antibodies.

### Western Blot

Protein extract was mixed with 2X XT buffer (Bio-Rad), boiled for 10 minutes, and resolved on a 4-12% gradient Bis-Tris gel (Bio-Rad). Proteins are on the gel transferred to the PVDF membrane, and specific antibodies were used to detect proteins of interest.

### RNA-seq

Total RNA (DNA free) was isolated using RNeasy Mini Kit (Qiagen), and ribosomal RNAs were depleted using ribosomal RNA depletion kit (rRNA Depletion Kit, NEB). Ribosomal depleted RNA was used to make the RNAseq libraries using NEBNext Ultra Directional RNA Library Prep Kit for Illumina (NEB). All the libraries were checked through a Bio-analyzer for quality control purpose. Paired-End (PE) 150bp reads were generated using a HiSeq-4000 sequencer (Illumina). RNAseq were performed in triplicates.

### Generation of mESC *KO* lines using CRISPR-Cas9

Paired sgRNAs were designed to delete coding exons (exons 1-2) of *Kdm1a* and *Kdm6a.* These created 17kb and 2kb genomic deletions of *Kdm1a* and *Kdm6a.* The sgRNAs were cloned into lentiguide-Puro (Addgene ID: 52963) plasmid, using Golden Gate Cloning approach as mentioned previously. Wild-type (J1) mESCs were transduced with Cas9-blast virus (generated from pLentiCas9-Blast, Addgene ID: 52962) and selected with 10μg/ml blasticidin (InvivoGen) to generate the J1 mESC cell line with stable expression of Cas9 (mESC+Cas9). 50,000 of mESC+Cas9 cells were transfected with 500ng of each sgRNAs (5 and 3’ sgRNAs) using Lipofectamine 2000 (Thermo Fisher Scientific), and cells were selected with 1μg/ml puromycin (Sigma-Aldrich) for 3-4 days. Next, puromycin-resistant cells were re-plated on a 15cm plate as single cells to grow individual clones. The individual clones were picked, expanded, and genotype PCR was performed to screen the homozygous/biallelic deletion or *knockout (KO)* mESC clones. Furthermore, Sanger sequencing and Western blot analysis were performed to validate *Kdm1a* and *Kdm6a KOs*.

### ChIP-seq analysis

The 50bp Single-End (SE) or 150bp Paired-End (PE) reads were mapped to the mm9 mouse genome assembly using Bowtie 2 (v2.3.2) ^40^. BigWig tracks were generated using the ‘bamCoverage’ program from the DeepTools software suite (v2.5.3) ^41^, with reads extended to 300bp and tracks normalized using the RPKM (Reads Per Kilobase per Million mapped reads) algorithm. Peak calling was done with the MACS2 program (v2.1.1.20160309) ^42^. For most ChIP-seq datasets default MACS2 settings were used. Artefact peaks were filtered out based on the mouse ENCODE project ^43^; they were removed using “bedtools intersect” from the BEDTools software suite (v2.26.0) ^44^. Mapping of peaks to the nearest gene was done using “bedtools closest”. Mouse ESC-specific enhancers taken from the literature ^21^. Promoters defined as a +/-2kb of TSS.

### Heat-map

ChIP-seq datasets were generated using the “computeMatrix” and “plotHeatmap” programs of the DeepTools software suite (v.2.5.3) ^41^ for ChIP-seq heatmaps and profile plots of histone marks and transcription factors binding. Regions from −2kb to +2kb around the centre of the peaks were used, split into 10 or 50bp bins, with each bin receiving the mean of the BigWig signal scores across it.

### GO (Gene Ontology)

Gene Ontology ^45^ analysis was performed using the NIH DAVID website (v6.8) ^46^; default settings and the whole genome were used as background. Genes were identified using their Ensembl Gene IDs for the DAVID analysis, and their associated gene names were obtained from the GENCODE vM1 mouse transcriptome annotation and added to the resulting GO term enrichment tables.

### MOTIF analysis

Enriched motifs of transcription factors (TFs) were identified at the ChIP peaks target regions using the “Haystack” bioinformatics pipeline, as described previously ^47^. Each enriched motif assigned with motif logos, p-values (calculated with the Fisher’s exact test) and q-values, the central enrichment score, the average profile in the target regions containing the motif, and the closest genes for each region.

### Module analysis

The entire mouse genome (mm9) was divided into constant 500bp bin windows. The bins that are overlapped with enhancers and promoters were selected for enhancer and promoter modules analysis, respectively. The average signal of each of these bins for each track was used to generate the matrix file using deeptools. K-mean clustering (K=3) was performed using average signals in each of the obtained clusters (i.e. centroids) based on Z-scores. Module heat maps were generated by using pheatmap package in R. For enhancer module analysis; enhancer bins overlapping with annotated promoters were excluded.

### Network analysis

Binding sites of each HDMs were mapped to the closest regions of defining networks (PRC, CORE, and MYC), as well as to the closest genes. We drew two diagrams, one representing all the mappings between HDMs and network regions associated with promoters; another one representing all the mappings between HDMs and network regions associated with enhancers. Edges were coloured based on the designation of the mapped networks (PRC = red, CORE = blue, MYC = orange). Tick marks next to network regions indicate that differential expression of HDM bound network genes in the differentiated state compared to the undifferentiated mESCs (p<0.05 and absolute log2-fold-change>1).

### RNA-seq analysis

Reads were mapped to the mm9 mouse genome assembly and the GENCODE version M1 mouse transcriptome ^48^ using the STAR RNA-seq read alignment program (v2.5.3a) ^49^ with default parameters. Bigwig tracks were generated using STAR to generate RPM-normalized wiggle files. Subsequently, the wigToBigWig tool (v4) from the UCSC genome browser Kent tools ^50^ was used to convert the wiggle files to BigWig format. Gene expression quantification, both raw read counts and TPM (Transcripts Per Million) values were obtained using htseq-count, part of the HTSeq package. Differential gene expression analysis was done using the DESeq2 (v1.14.1) ^51^ R statistical programming language (v3.3.2) software package from the Bioconductor project (v3.4) ^52^ using the GENCODE version M1 mouse transcriptome. DESeq2 was also used to calculate the normalized counts and the FPKM (Fragments Per Kilobase per Million mapped reads) expression values. MA plots were created employing the ggplot2 R package (v2.2.1) that used log2 fold changes and mean normalized counts calculated by the DESeq2 R package.

### Violin Plots/ differential expression

Log2 fold change for closest genes was plotted as violin plots using the **ggpbur R package (v0.4.0),** and the statistical significance of differential expression was calculated using the Wilcoxon rank-sum test with continuity correction, as implemented in the R statistical program (v3.3.2)

### Statistical analysis

All statistical analyses for the RNA-seq and ChIP-seq data were performed using the R statistical program (v3.3.2).

### Data availability

All the NGS data from this study have been deposited in GEO under the accession number GSE156107.

ChIP-seq data of HDMs were used from: Whyte et al., 2012. GSE27844 (KDM1A); Nair et al., 2012. GSE18515 (KDM1A); Farcas et al., 2012. GSE41267 (KDM2A, KDM2B); Pedersen et al., 2016. GSE64254 (KDM4A); Das et al., 2014. GSE43231 (KDM4B, KDM4C); Pedersen et al., 2014. GSE53936 (KDM4C); Kidder et al., 2014. GSE53087 (KDM5B); Schmitz et al., 2011. GSE31968 (KDM5B); Outchkourov et al., 2013. GSE34975 (KDM5C); Wang et al., 2017. GSE97703 (KDM6A); Banaszynski et al., 2013. GSE42152 (KDM6A, KDM6B); Højfeldt et al., 2019. GSE127804 (JARID2); Healy et al., 2019. GSE127121 (JARID2).

ChIP-seq data and RNA-seq data were used from Das et al., 2014. GSE43231; Seruggia et al., 2019. GSE113335.

ChIP-seq data for modules and networks analyses were used from Das et al., 2014. GSE43231; Seruggia et al., 2019. GSE113335.

ChIP-seq and RNA-seq data of Kdm4c KD were used from Das et al., 2014. GSE43231.

ChIP-seq and RNA-seq data of Kdm5b KD were used from Kidder et al., 2014. GSE53093.

RNA-seq data of *Jarid2 KO* were used from Das et al., 2015. GSE58414.

RNA-seq data of undifferentiated mESCs (0hr) and differentiated state (96hr) were obtained from Canver et al., 2019. GSE140911.

### Software and Algorithms

**1) BEDTools (2.26.0):** ^44^

http://bedtools.readthedocs.io/en/latest/

**2) Bowtie 2 (2.3.2):** ^40^

http://bowtie-bio.sourceforge.net/bowtie2/index.shtml

**3) MACS2 (2.1.1.20160309):** ^42^

https://github.com/taoliu/MACS

**4) DeepTools (2.5.3):** ^41^

https://deeptools.readthedocs.io/en/develop/

**5) STAR (2.5.3a):** ^49^

https://github.com/alexdobin/STAR

**6) HTSeq package (0.12.4):** ^53^

https://github.com/htseq/htseq

**7) wigToBigWig, Kent tools (v4):** ^50^

http://hgdownload.cse.ucsc.edu/admin/exe/

**8) DESeq2 R package (1.14.1; Bioconductor 3.4):** ^51^

https://bioconductor.org/packages/release/bioc/html/DESeq2.html

**9) DiffBind R package (2.2.12; Bioconductor 3.4):** ^54^

https://bioconductor.org/packages/release/bioc/html/DiffBind.html

**10) R statistical program (3.3.2):**

R Core Team (2017). R: A language and environment for statistical computing. R Foundation for Statistical Computing, Vienna, Austria.

URL https://www.R-project.org/.

**11) ggplot2 R package (2.2.1):**

ggplot2: Elegant Graphics for Data Analysis.

Springer-Verlag New York, 2009. ISBN: 978-0-387-98140-6

http://ggplot2.org

**12) pheatmap R package (1.0.8):**

pheatmap: Pretty Heatmaps. R package version 1.0.8.

https://CRAN.R-project.org/package=pheatmap

**13) NIH DAVID website (6.8):** ^46^

https://david.ncifcrf.gov/

**14) ggpubr R package (0.4.0)**

‘ggplot2’: Based Publication Ready Plots https://rpkgs.datanovia.com/ggpubr/

**15) Haystack:** ^47^

https://github.com/lucapinello/Haystack

## Acknowledgments

We thank Dr Wang and Prof Roeder for helpful discussions. We also thank Genewiz high-throughput sequencing facility. This work was supported by National Health and Medical Research Council (NHMRC) of Australia (APP1159461 and APP1182804) to P.P.D. L.P. is supported by a National Human Genome Research Institute (NHGRI) Career Development Award (R00HG008399), and Genomic Innovator Award (R35HG010717). S.H.O. is an Investigator of the Howard Hughes Medical Institute (HHMI).

## Author contributions

P.P.D conceptualised the study. Y.K., P.T., M.M., P.D., Z.Z., M.J.B., V.K.P., D.H., M.K., A.W., A.G., A.B.C and L.W. performed the experiments and analysed the data. Y.K., P.T., M.M., L.W., J.K., L.P., S.H.O. and P.P.D interpreted the data. Y.K., P.D., K.G., G.C.Y. and L.P. performed bioinformatics analyses. Y.K., P.T., L.P., S.H.O. and P.P.D. wrote the manuscript.

## Competing interests

The authors declare no competing interests.

## Notes

### Competing Interest Statement

The authors have declared no competing interest.

### Summary of Updates

- changes of author list - methods part

