## supplemental information for "Histone demethylome map reveals combinatorial gene regulatory functions in embryonic stem cells"

### Extended Data Fig. 1

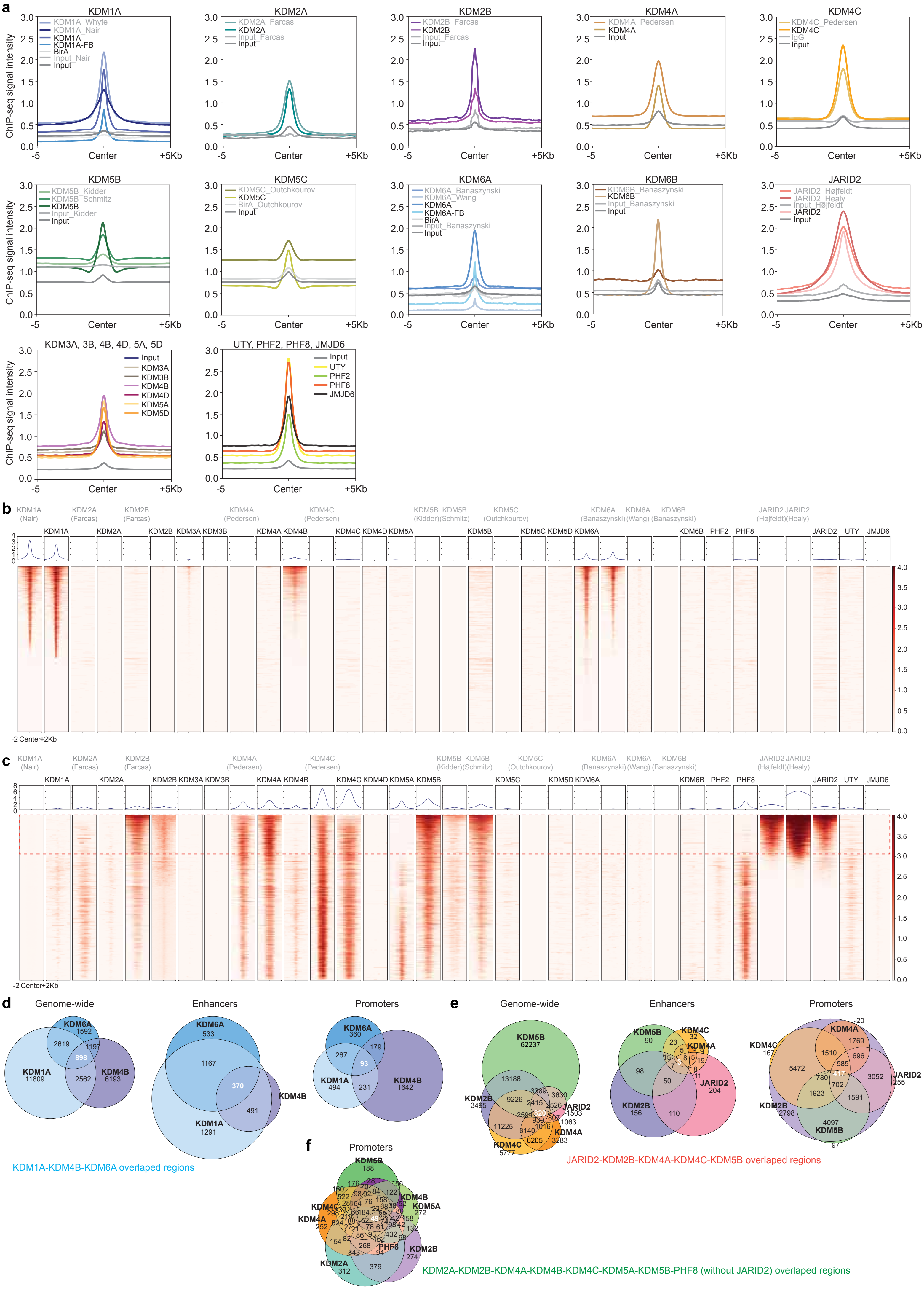

**Extended Data Fig. 2**

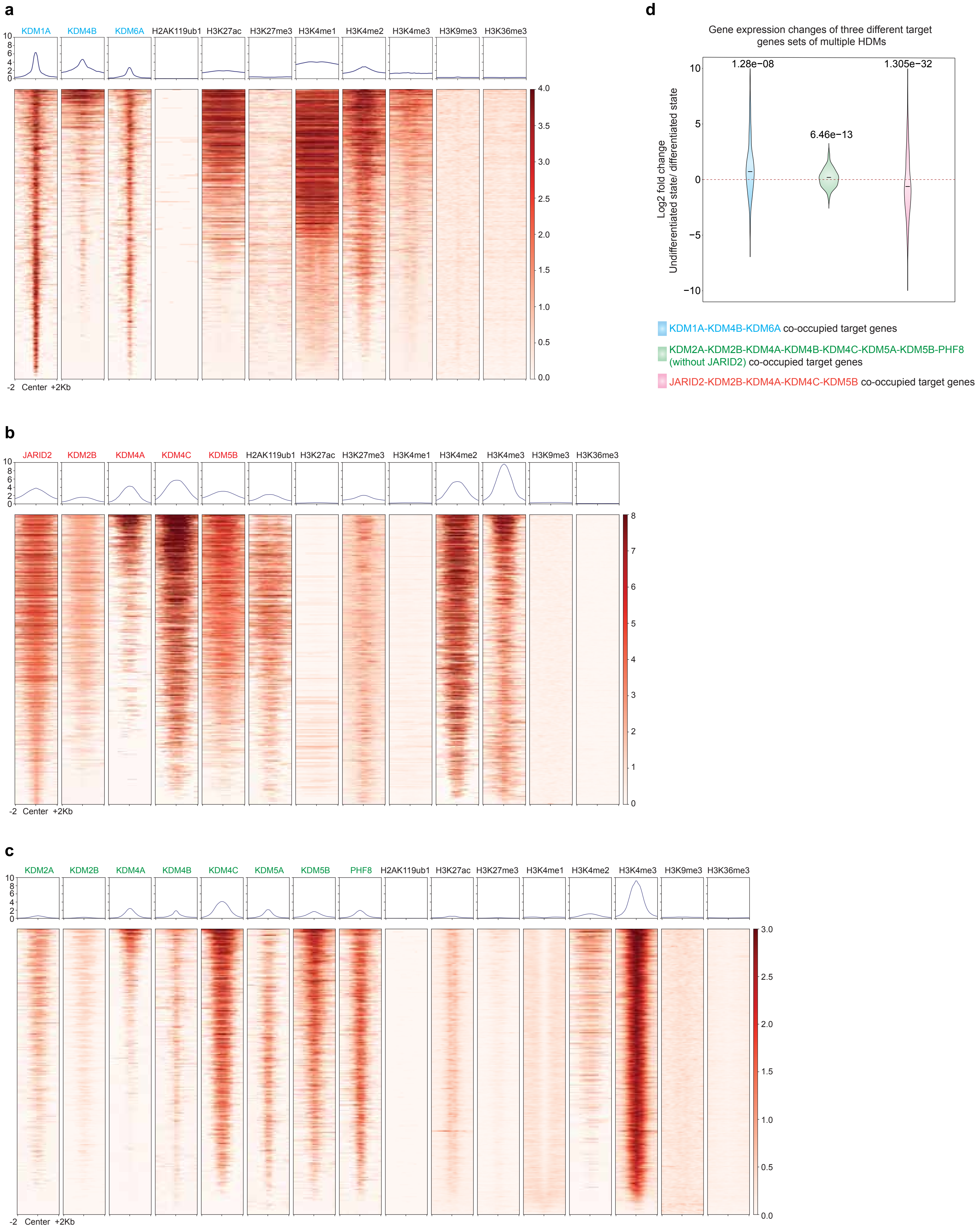

### Extended Data Fig. 3

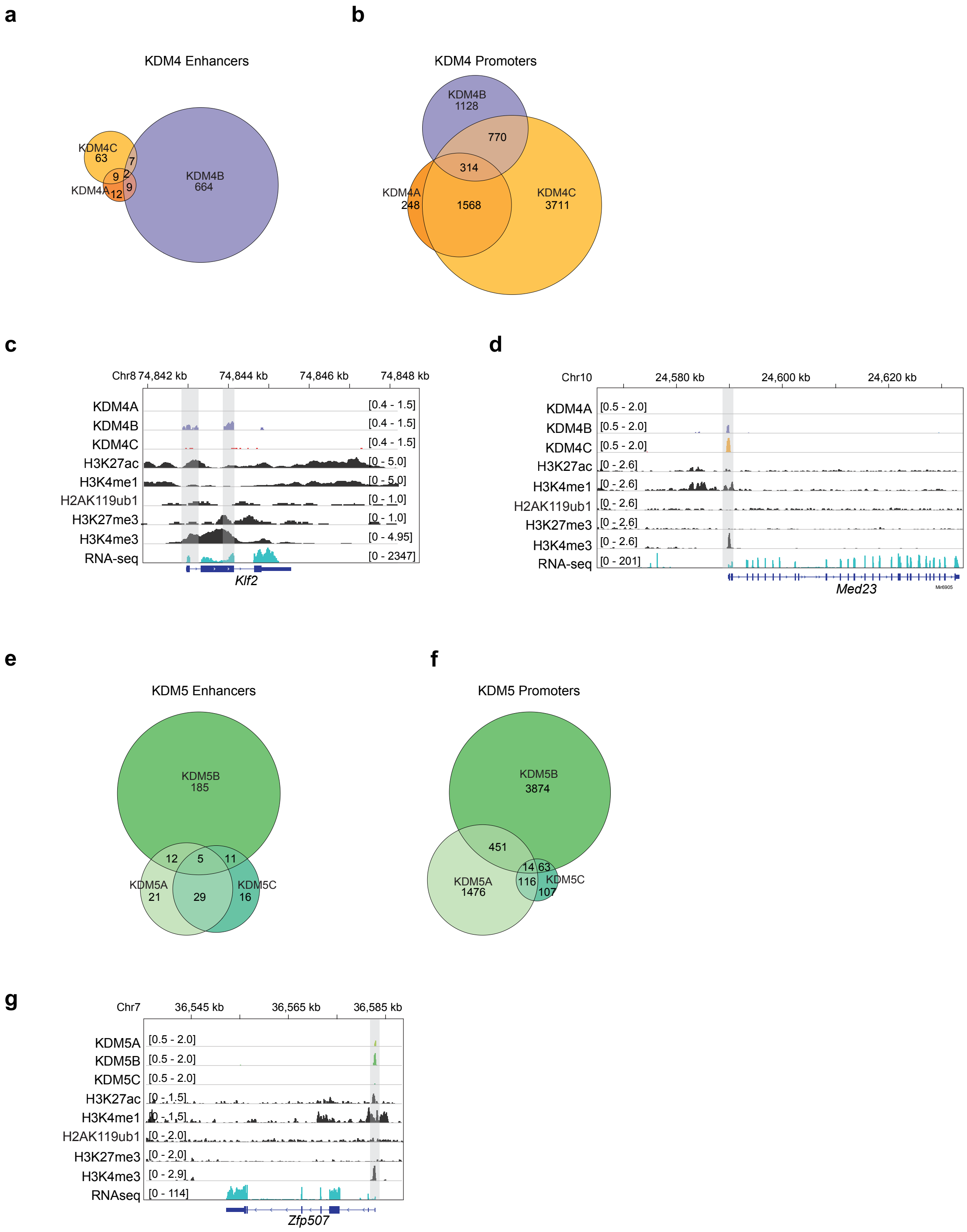

### Extended Data Fig. 4

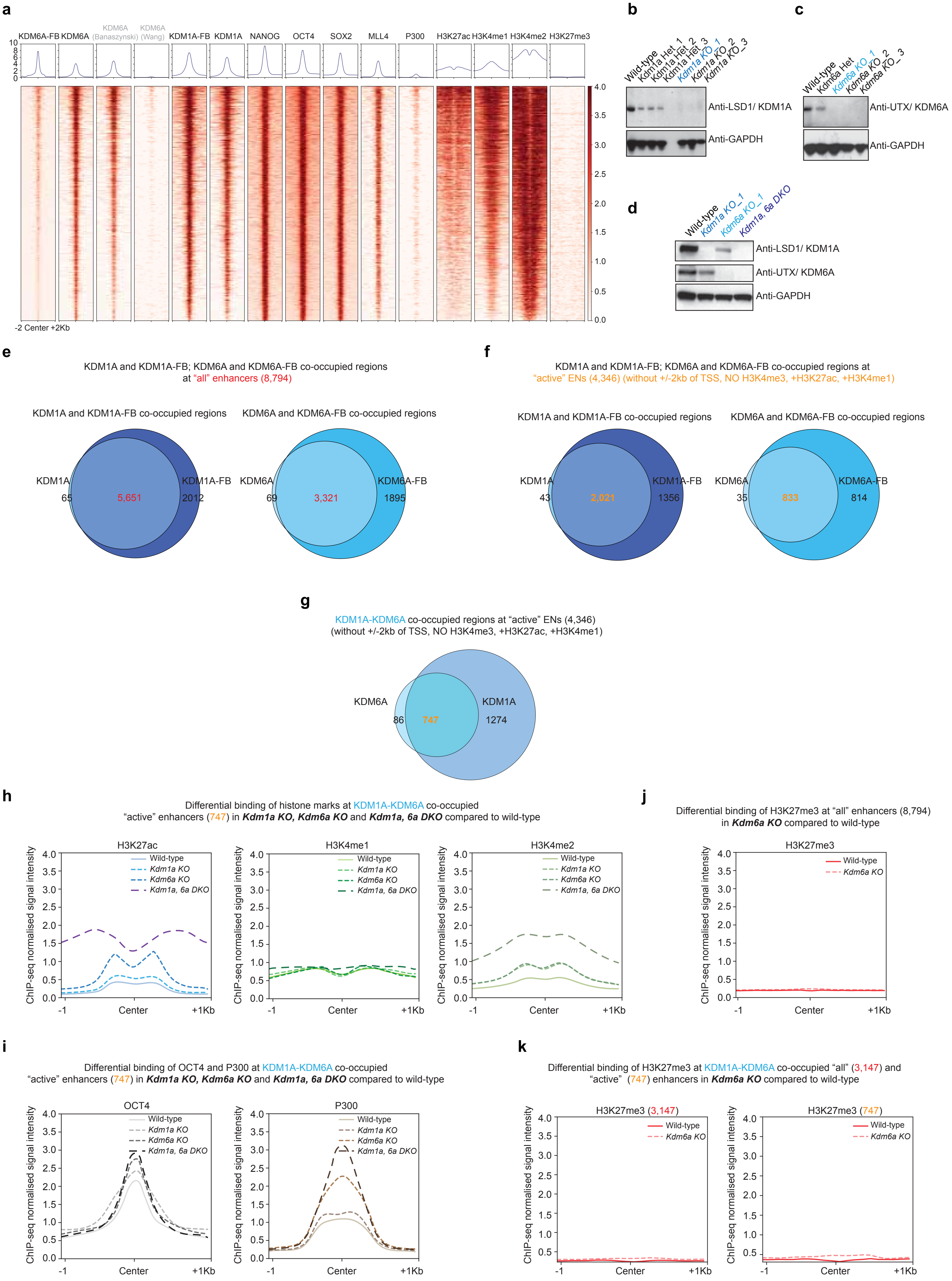

### Extended Data Fig. 5

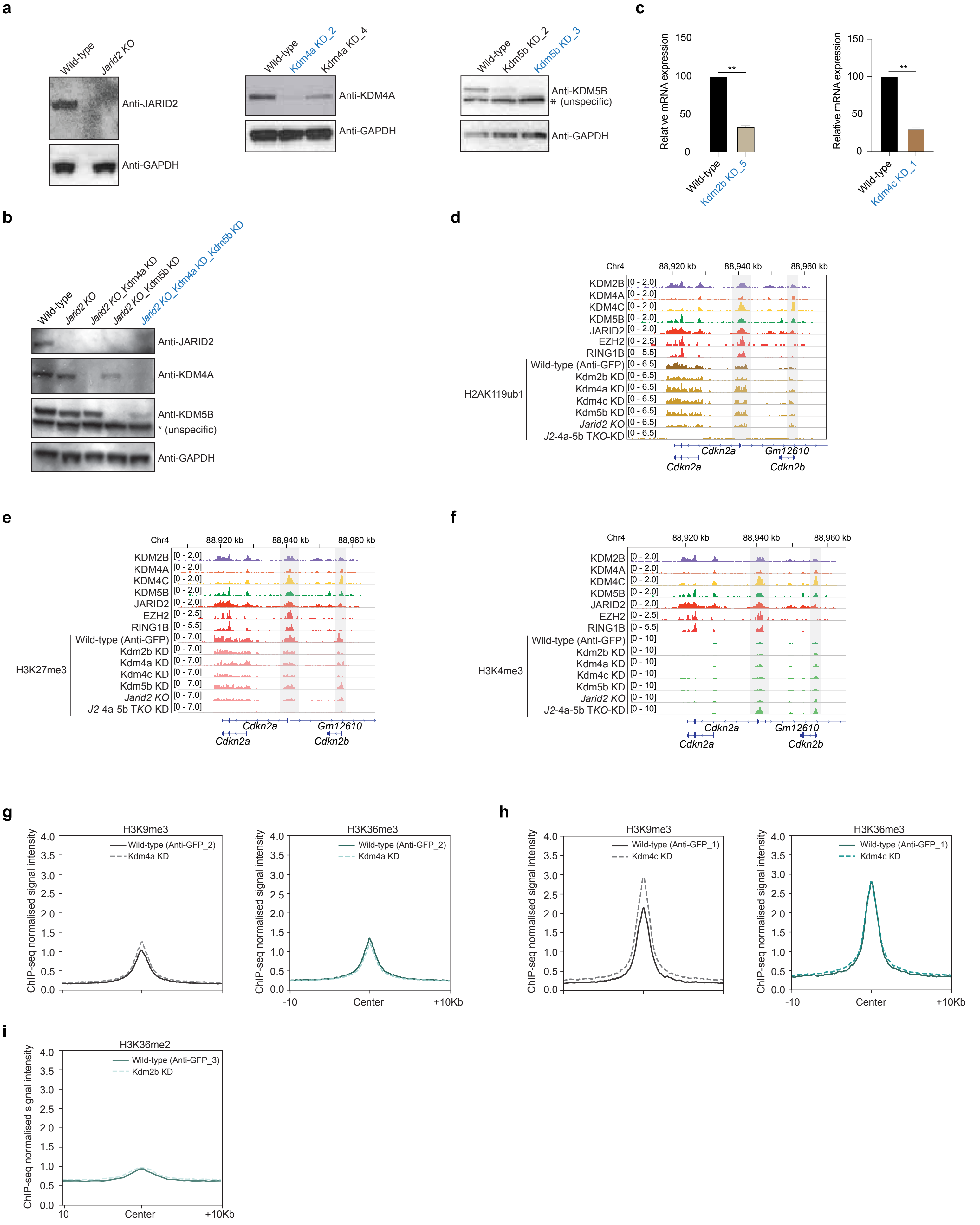

**Extended Data Fig 1. The HDMome map reveals the combinatorial co-occupancy of multiple HDMs.**

(a) Profile plots display ChIP-seq signal intensities of specific HDMs, as well as inputs/BirA (used as controls) in wild-types mESCs. Our ChIP-seq data (highlighted in black) and other published ChIP-seq data (highlighted in light grey, and mentioned with author names) of HDMs (KDM1A, KDM2A, KDM2B, KDM4A, KDM4C, KDM5B, KDM5C, KDM6A, KDM6B and JARID2) were compared from wild-type mESCs. In addition, our data represent ChIP-seq signal intensities of remaining HDMs – KDM3A, KDM3B, KDM4B, KDM4D, KDM5A, KDM5D, PHF2, PHF8, UTY and JMJD6 from wild-type mESCs.

(b, c) Heat maps represent co-occupancy of all HDMs at mESC-specific enhancers (b) and promoters (c). Plots are centred on the region midpoint ± 2kb. Relative ChIP-seq peak intensities are indicated. Our ChIP-seq data (highlighted in black) and published ChIP-seq data (highlighted in light grey, and mentioned with author names) of HDMs were used for comparison.

(d) Venn diagrams display overlapped binding regions of KDM1A, KDM4B and KDM6A at genome-wide, enhancer and promoter regions.

(e) Venn diagrams show overlapped binding regions of JARID2, KDM2B, KDM4A, KDM4C, KDM5B at genome-wide, enhancer and promoter regions.

(f) Venn diagram represents overlapped binding regions of KDM2A, KDM2B, KDM4A, KDM4B, KDM4C, KDM5A, KDM5B and PHF8 (without JARID2) at promoter regions.

**Extended Data Fig 2. Multiple HDMs combinatorially regulate gene expression.**

(a, b, c) Heat map represents co-occupancy of multiple HDMs and histone marks. Plots are centred on the region midpoint ± 2kb. Relative ChIP-seq peak intensities are indicated.

(d) Gene expression changes of three different target genes sets of multiple HDMs in the undifferentiated state compared to the differentiated state of mESCs.

**Extended Data Fig 3. Members of specific HDM sub-classes act combinatorially.**

(a, b) Venn diagrams show overlapped binding sites of KDM4A, KDM4B and KDM4C at enhancers (a) and promoters b).

(c, d) Genomic tracks represent ChIP-seq normalised reads for KDM4 members and histone marks at two different gene loci (*Klf2* and *Med23*). RNA-seq tracks demonstrate expression of these genes in wild-type mESCs.

(e, f) Venn diagrams display intersected binding sites of KDM5A, KDM5B and KDM5C at enhancers (e) and promoters (f).

(g) Genomic tracks represent ChIP-seq normalised reads for KDM5 members and histone marks at the *Zfp507* gene locus. RNA-seq track demonstrates expression of this gene in wild-type mESCs.

**Extended Data Fig 4. KDM1A and KDM6A combinatorially modulate P300/H3K27ac, H3K4me2 deposition and OCT4 recruitment at enhancers for target gene expression.**

(a) Heat map represents co-occupancy of KDM1A, KDM1A-FB, KDM6A, KDM6A-FB, ESC-TFs (OCT4, NANOG, SOX2), MLL4, P300 and histone marks (H3K7ac, H3K4me1, H3K4me2 and H3K27me3). Our KDM6A, KDM6A-FB ChIP-seq data and published KDM6A ChIP-seq data (Banaszynski and Wang) were used for the comparison.

(b, c, d) Western blot analysis confirmed *Kdm1a KO, Kdm6a KO* and *Kdm1a, 6a DKO* mESC lines. Highlighted *KO* mESC lines (blue in colour) were used for further experiments.

(e) Venn diagrams show KDM1A and KDM1A-FB co-occupied sites (5,651); KDM6A and KDM6A-FB co-occupied sites (3,321) at “all” enhancers (8,794).

(f) “Active” enhancers defined as: without +/- 2kb of TSS, NO H3K4me3, + H3K7ac, + H3K4me1. Venn diagrams display KDM1A and KDM1A-FB co-occupied regions (2,021); KDM6A and KDM6A-FB co-occupied regions (833) at active enhancers (4,346). The active enhancers (4,346) represent as a sub-set of all enhancers (8,794).

(g) Venn diagram shows high-confidence “KDM1A-KDM6A co-occupied regions” (747) at active enhancers (4,346). KDM1A and KDM1A-FB co-occupied sites (2,021), and KDM6A and KDM6A-FB (833) co-occupied sites at active enhancers were used to evaluate KDM1A-KDM6A overlapping binding sites (747) (see Extended Data Figure 5f).

(h, i) Profile plots represent differential binding of H3K7ac, H3K4me1, H3K4me2 (h), and OCT4, P300 (i) at KDM1A-KDM6A co-occupied active enhancers (747) in the *Kdm1a KO, Kdm6a KO* and *Kdm1a, 6a DKO* compared to wild-type.

(j) Profile plot show differential binding of H3K7me3 at all enhancers (8,794) in *Kdm6a KO* compared to wild-type.

(k) Profile plots display differential binding of H3K7me3 at KDM1A-KDM6A co-occupied enhancers (3,147 and 747) in *Kdm6a KO* compared to wild-type.

**Extended Data Fig 5. JARID2, KDM2B, KDM4A, KDM4C and KDM5B co-operatively control H2AK119ub1 (of PRC1) and bivalent marks (of PRC2) at the promoters for target gene repression.**

(a, b) Western blot analysis confirmed individual *Jarid2 KO (Knockout),* Kdm4a KD (Knockdown), Kdm5b KD (Knockdown) (a); as well as combined *Jarid2 KO*-Kdm4a KD-Kdm5b KD triple *KO*-KD (*J2*-4a-5b T*KO*-KD) (b) mESC lines. The highlighted mESC lines (in blue) were used for further experiments.

(c) Quantitative RT-PCR analysis confirmed Kdm2b KD and Kdm4c KD mESCs. Relative mRNA expression levels of Kdm2b and Kdm4c were shown in their respective KD mESC lines compared to wild-type. Data are represented as mean ± SEM (n=3); p-values were calculated using t-test. **p<0.005.

(d, e, f) Genomic tracks demonstrate differential binding of histone marks (H2AK119ub1, H3K27me3 and H3K4me3) at JARID2-KDM2B-KDM4A-KDM4C-KDM5B co-occupied promoters of the *Cdkn2a, 2b* gene loci in *KO* and KD mESCs compared to wild-type. The highlighted regions display differential binding of histone marks (*KO* and KD v. wild-type) at the JARID2-KDM2B-KDM4A-KDM4C-KDM5B co-occupied promoters.

(g, h) Profile plots represent differential binding of H3K9me3 and H3K36me3 at JARID2-KDM2B-KDM4A-KDM4C-KDM5B co-occupied promoters in the Kdm4a KD (g) and Kdm4c KD (h) compared to wild-type.

(i) Profile plot represents differential binding of H3K36me2 at JARID2-KDM2B-KDM4A-KDM4C-KDM5B co-occupied promoters in the Kdm2b KD compared to wild-type.
